# High-resolution mapping of the period landscape reveals polymorphism in cell cycle frequency tuning

**DOI:** 10.1101/2021.05.10.442602

**Authors:** Zhengda Li, Shiyuan Wang, Meng Sun, Minjun Jin, Daniel Khain, Qiong Yang

**Author notes:** These authors contributed equally to this work (Zhengda Li, Shiyuan Wang, Meng Sun).

## Abstract

Many biological oscillators exhibit widely tunable frequency in adapting to environmental changes. Although theoretical studies have proposed positive feedback as a mechanism underlying an oscillator’s large tunability, there have been no experiments to test it. Here, applying droplet microfluidics, we created a population of synthetic cells, each containing a cell-cycle oscillator and varying concentrations of cyclin B mRNAs for speed-tuning and positive-feedback inhibitors for modulating network interactions, allowing a continuous mapping of the cell-cycle period landscape in response to network perturbation. We found that although the cell cycle’s high tunability to cyclin B can reduce with Wee1 inhibition, the reduction is not as great as theoretically predicted, and another positive-feedback regulator, PP2A, may provide additional machinery to ensure the robustness of cell cycle period tunability. Remarkably, we discovered polymorphic responses of cell cycles to the PP2A inhibition. Droplet cells display a monomodal distribution of oscillations peaking at either low or high PP2A activity or a bimodal distribution with both low and high PP2A peaks. We explain such polymorphism by a model of two interlinked bistable switches of Cdk1 and PP2A where cell cycles exhibit two different oscillatory modes in the absence or presence of PP2A bistability.

## Introduction

The period of a cell cycle ranges broadly across different cells and organisms from as short as 8 minutes in Drosophila embryonic cells (Lehner & O’Farrell, 1989) to 24 hours in adult mammalian cells (Goldbeter & Berridge, 1996). The cell cycle can also adjust its period in a wide dynamic range in response to the changes of various intracellular and extracellular environmental conditions, including temperature (Francis & Barlow, 1988), nutrition availability (Cai & Tu, 2012), as well as the nuclear-cytoplasmic ratio that has been associated with the cell cycle period lengthening at the midblastula transition in early embryos (Levy & Heald, 2010). This ability to change the period following environmental perturbations without significant impact on the amplitude is known as tunability in frequency, critical to many biological oscillators.

Theoretical studies on the topological designs of biological oscillators (Tsai et al, 2008; Woods et al, 2016) have suggested that the positive feedback loops may play an important role in the robustness and great tunability of an oscillator. Modeling of the cell cycle network has also highlighted the essential role of Cdk1/Wee1/Cdc25 positive feedback in the cell cycle behavior as a relaxation oscillator, which provided an explanation why the cell cycle is highly tunable in frequency (Tsai et al, 2008; Gérard et al, 2012). However, there has been no direct experimental evidence to date to support this theoretical hypothesis. In addition, recent research shows that Wee1/Cdc25-based positive feedback may not be necessary for the cell cycle to oscillate, and positive feedback from phosphatase, especially PP2A, may also contribute significantly to cell cycle regulations (Krasinska et al, 2011; Mochida et al, 2016; Kamenz et al, 2021). These raise the questions of whether and to what degree each of the known Cdk1 positive feedback loops contributes to the regulation of the cell cycle period.

In a live adult cell, due to various checkpoints and interactions with other complicated cellular pathways, it is difficult to systematically modulate the strength of the positive feedback and investigate the response of the cell cycle as a free-running oscillator. Other major experimental challenges of studying cell-cycle tunability are the requirements for multi-dimensional network perturbation, high-resolution tuning, high-throughput sample preparation, and long-term single-cell imaging and analysis.

To overcome these challenges, we encapsulate *Xenopus* egg extracts that can cycle in the absence of checkpoints into cell-sized microfluidic droplets (Guan et al, 2018a, 2018b; Sun et al, 2019). To enable a multi-dimension, high-throughput, high-resolution mapping of the period landscape, we developed a programmed pressure-driven multi-channel microfluidic tuning system to gradually adjust ratios of cytoplasmic extracts containing the tuning drugs or mRNAs in individual droplets, resulting in a continuous spectrum of titration of these variables. Using this system, we found that the cell cycle period is highly tunable to cyclin B1 mRNA abundance, and such tunability can reduce when Wee1 activity is fully repressed, suggesting the Wee1-driven feedback indeed helps the cell cycle period tunability. However, its reduction extent is limited as compared to the model prediction, indicating a possibility of other redundant machinery that helps maintain the period tunability. We next perturbed PP2A, another positive feedback regulator of Cdk1, which like Wee1, affects the mitotic entry, but also controls the mitotic exit. We found that the cell cycle exhibits three qualitatively different behaviors concerning both oscillation probability and period tuning in response to PP2A inhibition. We build a cell cycle model to explain these observations to provide mechanistic insights into the role of the feedback regulations in cell cycle tunability.

## Results

### A high-throughput microfluidic platform to study *Xenopus* embryonic cell cycle period tunability

To quantitatively investigate the cell cycle tunability, we developed a platform that supports high-dimensional, high-resolution, high-throughput tuning of the cell cycle. We used the well-studied *Xenopus* egg extract system to reconstitute the early embryonic cell cycle (Fig 1A) since it allows us to focus on the nature of the mitotic circuit without other interfering gatekeeping mechanisms. We manufactured microfluidic devices to encapsulate cycling egg extract in cell-sized droplets with single-layer surfactant membranes. These droplets will act as artificial cells that can go through cell cycles without the complication of cell growth and cytokinesis. With this platform, thousands of artificial cells could be generated within minutes, followed by live imaging, automatic image processing, and statistical analysis.

**Figure 1.**
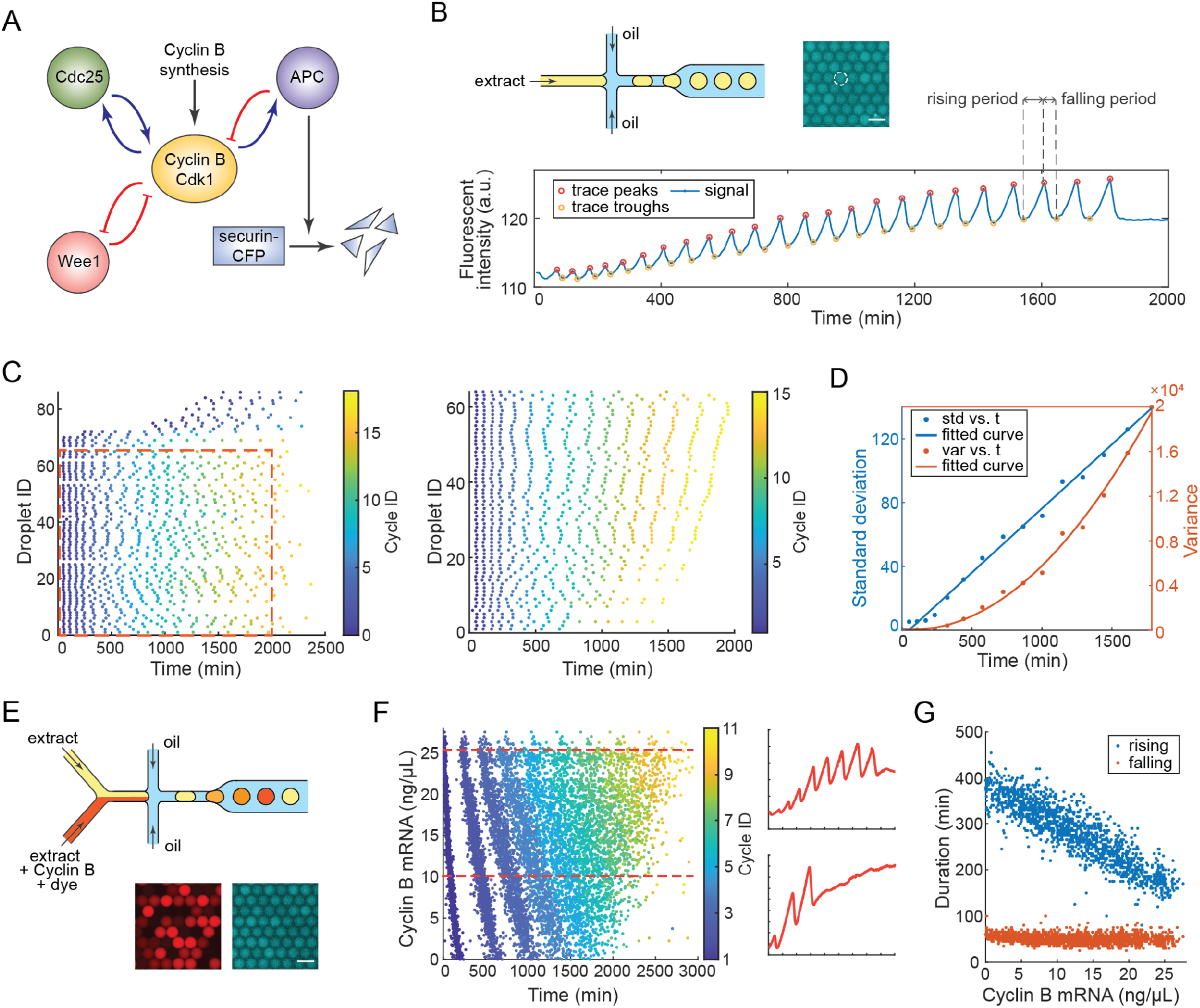
A high-throughput microfluidic platform to study *Xenopus* embryonic cell cycle period tunability. A Schematic view of the mitotic circuit with positive-plus-negative feedback loop design. Securin-CFP is used as the cell cycle indicator. B Top left: schematic view of a single-channel microfluidic device. Top right: a fluorescent image of droplets generated supplied with securin-CFP. Scale bar 100 µm. Bottom: the temporal fluorescent profile of an example droplet. The peaks and troughs are selected by custom MATLAB scripts and then manually corrected. C Left: Raster plot of the mitotic oscillations over time. Each dot represents a single peak in the fluorescent profile of one droplet placed at the time of the peak. Color bar indicates cycle ID. Right: Raster plot of the same dataset after optimization. Incomplete tracks are discarded, the remaining droplets (the selected area from the left figure) are sorted by average period length and reassigned droplet IDs. All points beyond the 15th cycle are also discarded. D standard deviation and variance of peak times against time. Each point represents all droplets with the same cycle ID. E Top: schematic view of a two-channel microfluidic device. Fluorescent dye is co-added with cyclin B mRNA to index its concentration. Bottom: fluorescent images of droplets generated. Bottom Left: fluorescent dye. Bottom Right: securin-CFP. Scale bar 100 µm. F Left: Raster plot of droplets with varying concentrations of cyclin B mRNA. Color bar indicates cycle ID. Right: fluorescent profiles of two example droplets with different cyclin B mRNA concentrations, indicated by dashed lines in the left figure. G The rising and falling period vs cyclin B mRNA concentration. The definitions of the rising and falling phases of the fluorescent profile are indicated in figure 1B. Only periods from the first cycle are included in the figure for clarity.

For a basic proof-of-concept experiment, we used a single-channel microfluidic device to generate uniform droplets without any form of content manipulation (Fig 1B). A chimeric fluorescent protein securin-CFP mRNA is added to visualize cell-cycle dynamics. As a substrate of anaphase-promoting complex or cyclosome (APC/C), securin is synthesized throughout interphase until anaphase onset, when APC/C is activated and ubiquitinates the protein securin for degradation (Fig 1A). Each droplet showed an oscillating CFP signal, indicating a successful constitution of cell cycle activities (Fig 1B).

Initial analysis of the temporal fluorescent signal of securin-CFP in all individual droplets gives highly synchronized oscillations among the droplets for their beginning cycles, as shown in the raster time series of oscillation peaks (Fig 1C, left). But this synchronicity was gradually lost in later cycles. To identify the source of desynchronization, we plotted the peak time standard deviation and variance versus the time of peaks (Fig 1D). If the periods are highly uniform and the desynchronization is caused by random phase drift, the peak time variance should be linearly correlated with time (Cao et al, 2015). On the other hand, if the period difference among droplets is dominant, the standard deviation should be linearly correlated with time. Our result shows a clear linear relationship between the standard deviation and time (Fig 1D), indicating that there might be significant differences in oscillation periods of different droplets, even though all droplets come from the same batch of extract. The period differences among droplets also become notable by sorting all droplets according to their average period over the total number of cycles (Fig 1C, right). The mechanism that causes these period differences is unknown, but a possible source of the period variability among our droplets is the partitioning error generated during the microemulsion encapsulation process (Weitz et al, 2014).

The fluorescent profiles also show that the oscillations eventually slow down, with gradually increasing periods (Fig 1B, S1A), possibly due to the consumption of energy (Guan et al, 2018a). Considering the droplets are isolated by oil and have minimal exchanges with the environment, we only use the first three cycles for further analysis in this study as they more accurately represent the physiological intracellular environment.

To tune the cyclin B production rate in droplets, we applied a 2-channel microfluidic device (Fig1E, Fig S1B) and injected the cycling extract to one channel and extract supplied with cyclin B mRNA and a dye to the other, both with the same amount of securin-CFP as a cell-cycle reporter. By varying the pressure of the two inlets following a pre-programmed pressure profile (Fig S1C), we generated droplets with a wide and continuous range of cyclin B mRNA concentration, calibrated by the dye fluorescence intensity (Fig S1D). In Fig 1F, we show that the oscillation behavior of droplets changed significantly with the cyclin B mRNA concentrations. As the externally added cyclin B mRNA concentration increases from 0 to 25 ng/µL, the number of cycles increases from 4 to 9, while the oscillation period decreases monotonically from close to 500 min down to 200 min (Fig 1F), suggesting the cell cycle has a high tunability in frequency. However, not all phases of the cell cycle are equally affected. The rising phase of the securin-CFP time courses, corresponding to the start of interphase until onset of anaphase, decreases linearly, while the falling phase, corresponding to the start of anaphase until mitotic exit, largely remains constant (Fig 1G). The phase-dependent sensitivity of cell cycle to the cyclin B variation agrees with previous reports, where the mitotic phase is found to be temporally robust and insulated from other cell-cycle events that are more variable, a phenomenon postulated to be caused by the positive feedback loops in the mitotic circuit (Araujo et al, 2016).

### Inhibiting Wee1 positive feedback reduces period tunability

Next, we investigated the contribution of positive feedback loops to the large frequency tunability of the mitotic oscillator observed in our cyclin B mRNA tuning experiments (Fig 1E-G). Previous theoretical studies attributed the cell cycle tunability to the Cdk1/Wee1/Cdc25 positive feedback that couples with the time-delayed negative feedback loop formed between Cdk1 and APC/C (Tsai et al, 2008) (Fig 1A). The antagonistic pair of Wee1 kinase and Cdc25 phosphatase form two positive feedback loops with the master mitotic regulator cyclin B-Cdk1 complex, with active Cdk1 inhibiting its inhibitor Wee1 and activating its activator Cdc25.

We compromised the kinase activity of Wee1 using a tyrosine kinase inhibitor PD 166285 (Fig 2A). Since the antagonist Cdc25 removes the Wee1-mediated inhibitory phosphate groups of Tyr15 and Thr14 on Cdk1, the addition of PD 166285 can reduce the strength of both Wee1 and Cdc25 positive feedback loops. We first applied different amounts of PD 166285 manually to extracts before performing 2-channel cyclin B mRNA tuning to test if the frequency tunability observed in previous cyclin B tuning experiments will be affected. The results revealed that while droplets with no PD 166285 showed a wide range of oscillation frequencies, this range was dramatically reduced when the inhibitor of 2 µM or more was present (Fig 2B). In the group with the highest inhibitor concentration (10 µM), the period change is on the same level as background noise in periods. This result suggests that interrupting Wee1/Cdc25-based positive feedback did have an impact on cell cycle tunability.

**Figure 2.**
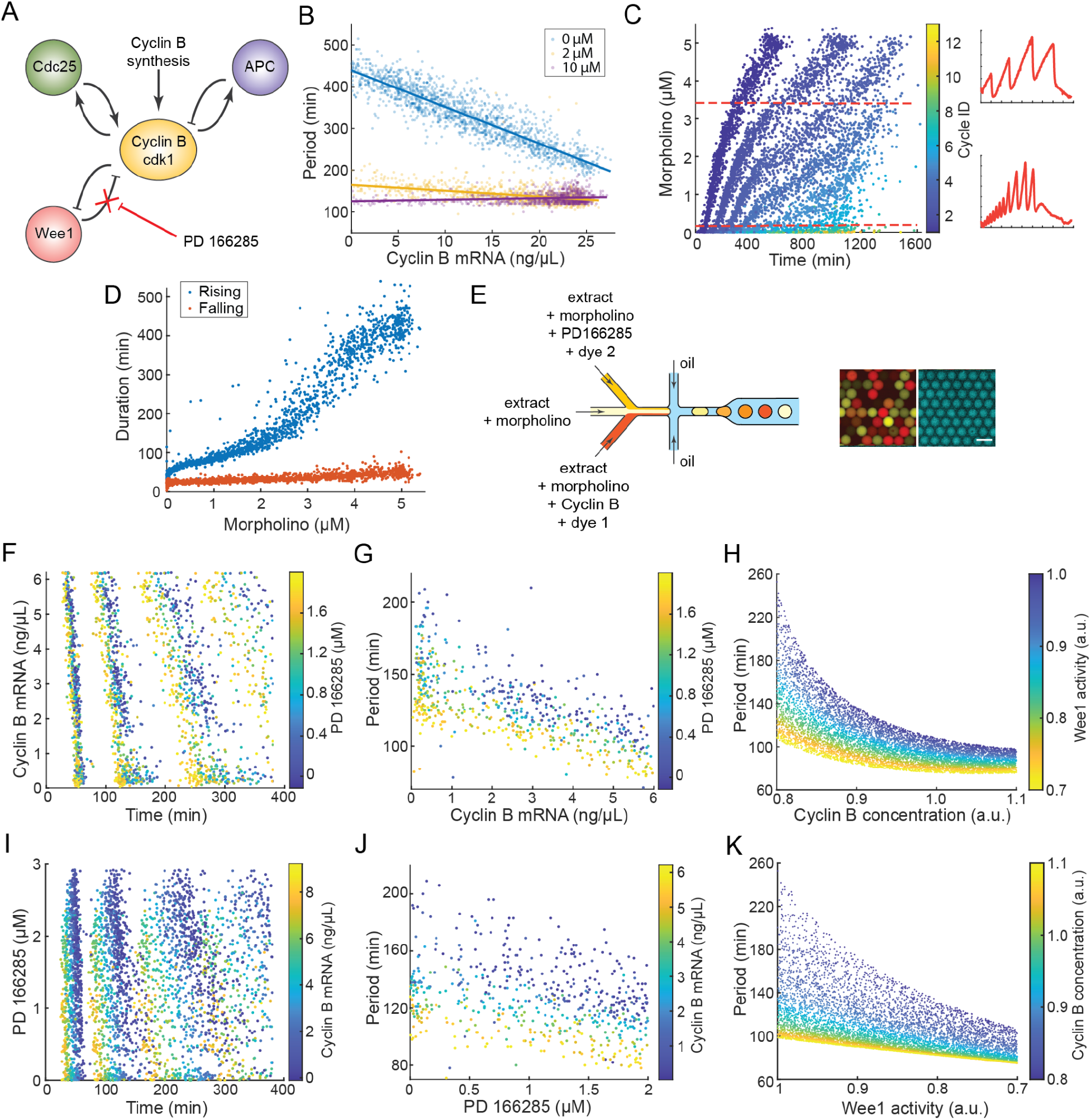
Inhibiting Wee1 positive feedback reduces period tunability. A Schematic view of the function of PD 166285 on the mitotic circuit. B Oscillation period vs. Cyclin B mRNA concentration. The differently colored scatters represent 3 distinct PD 166285 concentrations. The lines indicate linear approximations. Only periods from the first cycle are included in the figure for clarity. C Left: Raster plot of droplets with varying concentrations of morpholino. Color bar indicates cycle ID. Right: fluorescent profiles of two example droplets with different morpholino concentrations, indicated by dashed lines in the left figure. D The rising and falling period vs morpholino concentration. Only periods from the first cycle are included in the figure for clarity. E, F, G, I, J Results of three-channel tuning experiments. PD 166285 and cyclinB mRNA concentrations are tuned simultaneously. E Left: schematic view of a three-channel microfluidic device. 2 Fluorescent dyes are co-added with cyclin B mRNA and PD 166285 respectively to index their concentrations. Right: fluorescent images of droplets generated. Left: composite of fluorescent dyes. Right: securin-CFP. Scale bar 100 µm. F Raster plot of droplets with varying concentrations of cyclin B mRNA. Color bar indicates PD 166285 concentration. G Scatter of oscillation period vs Cyclin B mRNA concentration. Color bar indicates PD 166285 concentration. Only periods from the first cycle are included in the figure for clarity. H Simulated result of oscillation period vs Cyclin B activity. Color bar indicates Wee1 activity. I Raster plot of droplets with varying concentrations of PD 166285. Color bar indicates Cyclin B mRNA concentration. J Scatter of oscillation period vs PD 166285 concentration. Color bar indicates Cyclin B mRNA concentration. Only periods from the first cycle are included in the figure for clarity. K Simulated result of oscillation period vs Wee1 activity. Color bar indicates Cyclin B activity.

However, in these experiments, cyclin B tuning was on top of the endogenous cyclin B mRNAs already existed in *Xenopus* extracts, and the cell cycle could only go in the direction of increasing beyond its natural speed. How cells respond to the range of cyclin B lower than its endogenous level is missing. Moreover, the oscillation period also decreases moderately with the Wee1 activity inhibition alone, as observed in a 2-channel tuning experiment of PD 166285 (Fig S2A). Since both cyclin B mRNA and PD 166285 can speed up oscillations, when tuning in the range of high cyclin B mRNA concentrations, the addition of PD 166285 could drive the cell cycle period to a minimum where the system cannot be altered any further by cyclin B mRNA addition. This may provide an alternative explanation for the severely reduced sensitivity observed in Fig 2B.

To eliminate this possibility and map the behavior near the lower boundary of cyclin B, we applied a mixture of 4 morpholino antisense oligonucleotides to block the translation initiation of 4 endogenous *Xenopus* cyclin B (x-cyclin B) mRNA species, including 2 isoforms each of *c*yclin B1 and cyclin B2 (Materials and Methods). The addition of x-cyclin B morpholinos (MOs) allows slower endogenous oscillations and thus a wider dynamic range of tuning when we supply recombinant human cyclin B (h-cyclin B) mRNAs into these extracts. By 2-channel tuning of x-cyclin B MOs, we confirmed the effectiveness of the MOs to inhibit endogenous cyclin B expression (Fig 2C), with oscillations drastically slowing down as MO concentration increases. Consistent with the previous cyclin B tuning results, only the rising period changes significantly with morpholino concentration, while the falling period is almost unaffected (Fig 2D).

We also noticed that the coarse-grained manual tuning of Wee1 inhibition showed a sudden loss of tunability upon the addition of PD 166285, failing to capture the intermediate changes to Wee1 inhibition in between (Fig 2B). To systematically map how the system responds to Wee1 perturbation, we developed a 3-channel tuning device (Fig 2E, Fig S2E) to enable a fine-grained automatic tuning of both cyclin B mRNA and PD166285 simultaneously. For the aforementioned reason, we here focused on the low concentration regions of cyclin B and PD 166285 with a higher resolution. The cyclin B tuning is achieved by adding an excessive amount of morpholinos to silence all endogenous x-cyclin B mRNA and then adding human cyclin B mRNA to restore translation. We programmed a 3-inlet pressure profile (Fig S2F) so that the generated droplets have relatively uniform distributions of PD166285 and cyclin B mRNA concentrations (Fig S2G). The results showed that Wee1 activity still plays a role in the frequency tunability to cyclin B synthesis rate (Fig 2F, Fig 2G), but the effect is less obvious than the previous MO-untreated cyclin B tuning experiment. The difference is more evident in the lower cyclin B mRNA concentration region, where higher PD 166285 concentrations lead to significantly narrower ranges of periods to tune for the same span of cyclin B mRNA concentration. But for the higher cyclin B concentration region, changes in Wee1 activity have little effect on the oscillation period sensitivity to cyclin B.

As a comparison to the experimental observations, we revisited the theoretical hypothesis that the Wee1-based positive feedback of Cdk1 is necessary for the frequency tunability of cell cycle oscillators (Tsai et al, 2008) and applied a cell-cycle model (Yang & Ferrell, 2013) to calculate the period range as we tune cyclin B synthesis rate (Fig 2H) and Wee1 activity (Fig 2K) respectively. The model could capture the overall trends of the cell cycle period changes to cyclin B mRNA concentration at various Wee1 inhibition (Fig 2G vs Fig 2H) and to PD 166285 concentration at different cyclin B mRNA levels (Fig 2J vs Fig 2K). However, the theoretical result suggests that even at the low cyclin B concentration region (as in the morpholino pre-treated experiments), Wee1 inhibition should greatly reduce the cell-cycle period tunability to cyclin B mRNA, which is not as obvious in experiments. This discrepancy suggests that the tunability of the cell cycle is more robust than theoretically predicted, and there might be additional mechanisms other than Wee1/Cdc25-based positive feedback that can ensure a flexible oscillation period.

### Inhibiting PP2A leads to polymorphism in oscillation and period responses

Apart from the extensively characterized Cdk1/Wee1/Cdc25/ positive feedback, recent studies have identified additional positive feedback centered on the Cdk1-counteracting phosphatase, PP2A, formed by the mutual antagonism of the Endosulfine Alpha/Greatwall (ENSA/Gwl) pathway and PP2A (Mochida et al, 2016; Rata et al, 2018) (Fig 3A). Studies also suggested that PP2A contributes to the period regulation of the cell cycle (Naetar et al, 2014).

**Figure 3.**
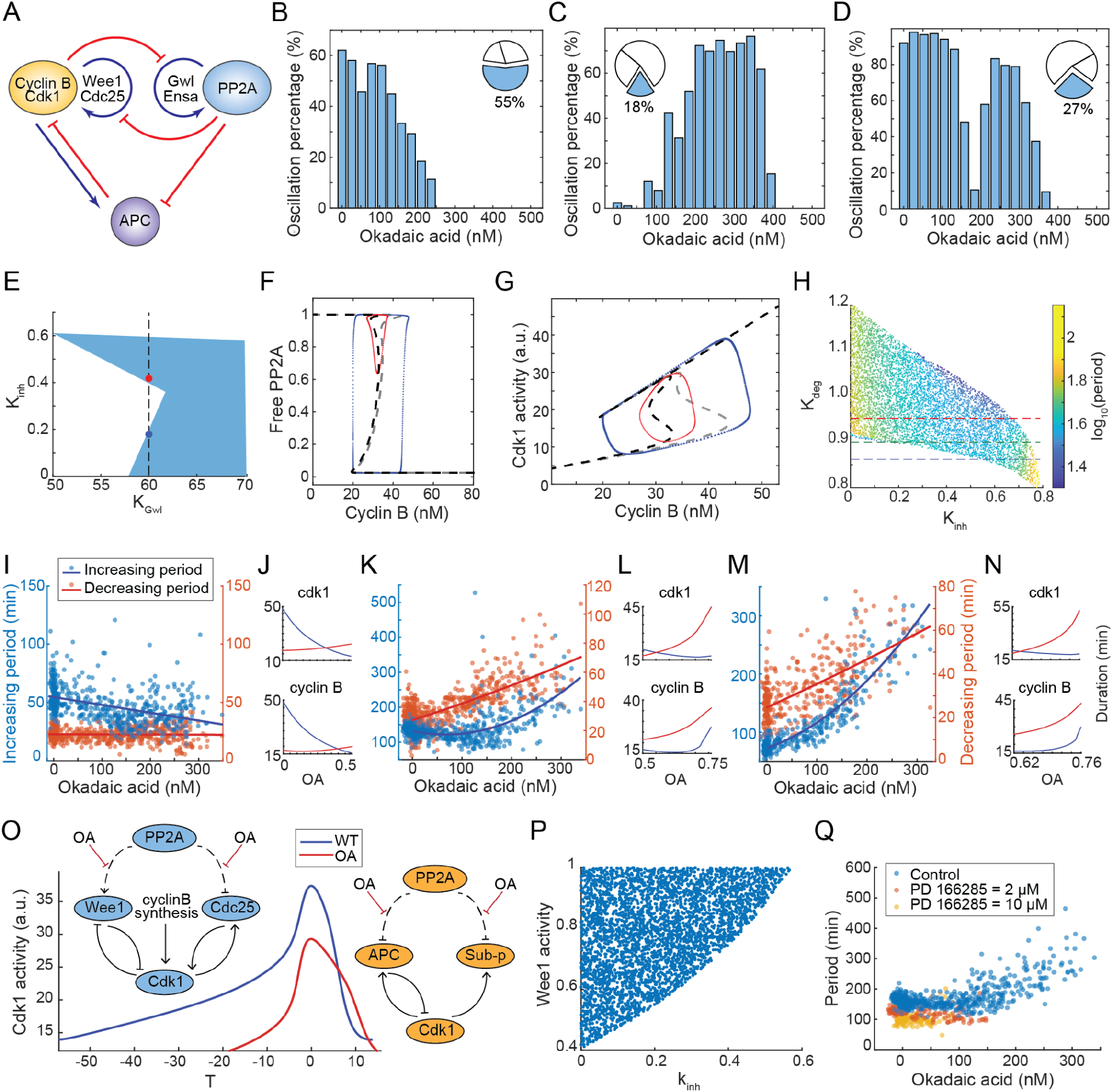
Inhibiting PP2A leads to polymorphism in oscillation and period responses. A Schematic view of molecular circuits including PP2A activities. B, C, D Percentage of oscillatory droplets vs OA concentration, showing type 1 (B) type 2 (C), and type 3 (D) responses. Inset: Pie plot shows the percentage of each type in all experiment results. E Parameter regions that support oscillation when changing K_Gwl_ and k_inh_ (OA), showing that changing k_inh_ at a specific value of K_Gwl_ (along the dotted black line) may lead to type 3 OA response. Blue and red dots show oscillations in low and medium OA concentration respectively. F, G Dashed lines show response curves of steady-state free PP2A (F) or Cdk1 (G) activity when changing Cyclin B concentration under low (gray) or medium OA (black). Dotted lines show phase plots of limit cycles with low (blue) or medium (red) OA corresponding to figure 3E. H Oscillation period (color coded) changes when tuning K_inh_ (OA concentration) and ec50_apc_ (activity threshold for Cdk1 substrate). Red, green, blue dotted lines show three different OA period responses. I, K, M Experimental results of increasing (blue) and decreasing (red) period of securin-CFP when tuning OA, showing three different types of OA period response: monotonically decreasing (I), decreasing and then increasing (K) or monotonically increasing (M). Solid lines are fitted curves with linear or quadratic regression. J, L, N Simulated results of increasing (blue) and decreasing (red) period of Cyclin B concentration (lower) and Cdk1 activity (upper) when adding OA, corresponding to three different types of OA period, showing that the period change of Cyclin B is similar to that of securin on the left, while the Cdk1 activity shows a similar changing pattern. O Representative time series of Cdk1 activity of WT (blue) and OA treated (red) cycles. The schematic molecular circuits show different OA effects during increasing and decreasing phases of oscillation. P Oscillatory region when tuning Wee1 activity and k_inh_ (OA) in a computational model, showing that Wee1 inhibition leads to a smaller region of oscillation when adding OA. Q Period responses to OA when adding PD 166285 in different concentrations.

To quantitatively understand the effect of PP2A on the cell cycle dynamics, we applied okadaic acid (OA), a classical PP2A inhibitor, in our droplet microfluidic tuning system to tune PP2A activity. Surprisingly, droplet cells respond to changes in OA concentration in a polymorphic manner, resulting in three distinct types of patterns of OA concentration among oscillatory cells. Here, we define oscillation percentage as the percentage of oscillatory cells among all recorded droplet cells within each bin of OA concentration. Type 1 shows a continuous decrease in oscillation percentage with increasing OA concentration (Fig 3B and S3A). Type 2 shows that the oscillation percentage initially increases, then decreases, and eventually drops down to 0 at a specific OA concentration (Fig 3C and S3B). Type 3 shows two OA concentration ranges supporting oscillation, separated by a gap where the cell cycle becomes arrested (Fig 3D and S3C). These observations are not due to the uneven distribution of OA concentrations in the entire population of droplets (Fig S3D-F). Out of 11 independent experiments, a majority showed a type 1 response (n=6), while type 2 (n=2 out of 11) and type 3 (n=3 out of 11) are rarer. The exact mechanism that determines the type of response is unclear, but it could be associated with different initial conditions of extracts caused by batch differences in eggs or day-to-day variability in extract preparation processes since droplets produced with the same bulk extracts from the same batch of eggs always show the same type of response.

Among all response types, type 3 displays a unique bimodal distribution that no studies have reported before. We could recapitulate this phenomenon by applying the alternative coarse-grained manual tuning method. We divided the same batch of extract into several groups each applied with different amounts of OA, and then with single-channel microfluidic devices, we generated droplets with uniform OA concentration for every group. For one experiment, the result clearly shows that while OA arrests oscillations for groups of 100 nM, 200 nM, and 800 nM, it supports oscillation at 0 nM, 50 nM, and 400 nM (Fig S3G).

To further understand the two-modal behavior of the type 3 response, we expanded our cell cycle model to include PP2A circuits based on a recent study (Kamenz et al, 2021) (Fig S3H). We explored a hypothesis that could explain the polymorphism by variations of certain parameters due to extract-to-extract variability. We find that type 3 oscillation could be achieved by changing K_Gwl_, the threshold of Cdk1 activation of Gwl, in our model to a certain value (Fig 3E). This threshold has been shown to vary among different frogs in previous experiments (Kamenz et al, 2021). To explain this behavior, we plotted the response curves of free PP2A and active Cdk1 to cyclin B (Fig 3F and 3G). Previous research had shown that the activation of Cdk1 by cyclin B is bistable (Pomerening et al, 2005), and the activation of PP2A by Cdk1 is also bistable (Mochida et al, 2016) (Fig S3I). Combining these two bistabilities, we could get a tri-stability response curve with regards to cyclin B. The existence of a third intermediate steady state in between the interphase and M phase of the cell cycle has also been predicted and experimentally verified in a previous study in mammalian cells (Rata et al, 2018).

Our model shows that the oscillation may occur among different steady states. When the OA level is low, the oscillation occurs between high and low steady states of PP2A, representing a full-amplitude cycle (Fig 3F and 3G, blue curve) when both Cdk1 bistability and PP2A bistability are present during oscillations. When the OA level is high, Wee1 and Cdc25 are hyperphosphorylated, making the Cdk1 positive feedback more sensitive. This leads to prematurely activated Cdk1. As a result, the peak Cdk1 level is too low to fully inhibit PP2A to trigger PP2A-ENSA-Gwl bistability. The oscillation occurs between the intermediate and high steady states of PP2A (Fig 3F and 3G, red curve), which represents a consistently high level of free PP2A. In this case, the oscillation is solely driven by the bistability based on Wee1/Cdc25 positive feedback.

This hypothesis, while not experimentally verified, could explain the polymorphism of oscillatory behaviors when tuning OA. The type 1 and the first mode of type 3 responses could be explained by a full-amplitude oscillation of PP2A as OA is low, while the type 2 and the second mode of type 3 responses could be explained by a partial-amplitude oscillation that would require partial PP2A inhibition.

In addition to oscillation percentage, the period of the oscillator also responds polymorphically to PP2A inhibition. Three types of responses have been observed, showing a monotonic decrease (Fig S3J), a monotonic increase (Fig S3L), and a slight decrease followed by an increase (Fig S3K). This phenomenon could be explained by a slight variation on K_deg_ in the model (Fig 3H), which represents the threshold for APC activation by Cdk1. To test this hypothesis, we measured the duration of increasing phase and decreasing phase in a cycle (Fig 3I, K, M) to compare with model predictions. Since cyclin B and securin are both APC substrates and share the same synthesis (increasing) and degradation (decreasing) phases, we could compare the increasing period and decreasing period experimentally measured by a securin-CFP reporter (Fig 3I, K, M) with those predicted from the simulated cyclin B dynamics (Fig 3J, L, N, lower panel). The results show high similarities between experimental and modeling results.

To further understand period polymorphism, we also plotted the changes in the increasing and decreasing periods of Cdk1 activity when tuning OA (Fig 3J, L, N, upper panel). Interestingly, while cyclin B’s response exhibits polymorphism, Cdk1’s response is conserved. In all three period-response types, the Cdk1’s increasing period always decreases with OA concentration while its decreasing period increases. This leads to a hypothesis that the period polymorphism is determined by the relative sensitivity of the increasing/decreasing phase of Cdk1 activity to OA. As illustrated in Fig 3O, OA can differentially affect the regulations of mitotic entry and mitotic exit. During mitotic entry, OA functions as an inhibitor of Wee1 and an activator of Cdc25, which reduces the threshold of Cdk1 activation and leads to accelerated cell cycle progression, while in mitotic exit, OA being an inhibitor of PP2A could delay cell cycle progression and even arrest oscillations. In addition, while fully inhibiting PP2A will cause cell cycle arrest, the PP2A bistability may not be necessary for cell cycle oscillations. The model predicts a moderate level of PP2A inhibition that could lead to prematurely activated Cdk1 and APC, but the Cdk1 level would be too low to drive PP2A bistability.

To understand how both the Cdk1/Wee1/Cdc25 and PP2A/ENSA/Gwl positive feedback loops work together to influence cell cycle period tuning, we simulated the oscillations in response to both dimensions of Wee1 inhibition as well as PP2A inhibition (Fig 3P) and found that inhibitions of PP2A and Wee1 can reduce each other’s ability to sustain oscillations. This result predicts that adding the Wee1 inhibitor can lead to a reduced OA range (k_inh_) that supports oscillations. To verify this prediction, we tuned the OA concentration in several groups of droplets, each with a distinct PD 166285 concentration. The result shows that higher PD 166285 indeed gives a more significantly reduced OA tuning range of oscillations (reduced robustness) and reduced period tunability (Fig 3Q).

## Discussion

Many biological oscillators have well-conserved positive feedbacks despite that, in theory, a single time-delayed negative feedback is sufficient to generate sustained oscillations.Although extensive computational work has been done suggesting that positive feedback may play a role in clock tunability and robustness, the essential experimental studies in this regard have been missing. We here developed a high-throughput experimental platform to simultaneously perturb positive feedback and map the period landscape of many droplet cells undergoing cell cycle progression. Instead of studying a few specific phenotypes, we analyzed the whole spectra of phenotypes by continuously varying parameters, which provides comprehensive information necessary for verifying theories and understanding the underlying mechanisms at the systems level.

We showed that the previously proposed Wee1/Cdc25 positive feedback only contributes partially to the tunability of the cell cycle. Since Wee1 inhibition itself can reduce the period by lowering the activation threshold of Cdk1, it interferes with the mRNA tuning especially towards the upper limit of the frequency range. As a result, adding excessive mRNA only contributes slightly to the period tuning when Wee1 inhibition is high, and morpholino is needed to search for the lower end of frequency under this condition. With the high-resolution mapping of the period landscape, our results revealed the resistance of tunability to perturbations in the Wee1 positive feedback, possibly due to the redundant positive feedback provided by PP2A. Previous studies also found that inhibition of Wee1 activity alone could not abolish the bistability of the cell cycle (Chang & Ferrell, 2013).

Different from Wee1 perturbations, the cell cycle response to PP2A inhibition is rather complicated. We have observed polymorphism in both oscillatory fractions and period tuning patterns. While we have not yet fully understood the mechanisms underlying these phenomena, our model suggests that the polymorphism could be explained by the relative sensitivity of Cdk1’s substrate Gwl or APC to Cdk1. The sensitivity of a substrate can be considered as to how soon the substrate is activated in response to Cdk1. This highlights the significance of the temporal order of activation of APC and Gwl, and a slight change in the temporal order of the cell cycle may lead to a dramatic change in oscillatory patterns.

In addition, while we do not know how much of our observations in egg extracts represent the physiological behavior of embryos, the results from droplet microfluidics mapping suggest that there might be more than one mode of cell cycle behaviors, and the oscillation properties and the temporal order of mitotic events might vary in these different oscillatory modes. These complex behaviors could not have been detected by low-throughput and low-resolution network perturbations in live embryos or bulk extract assays.

## Materials and Methods

### Reagents and Tools Table

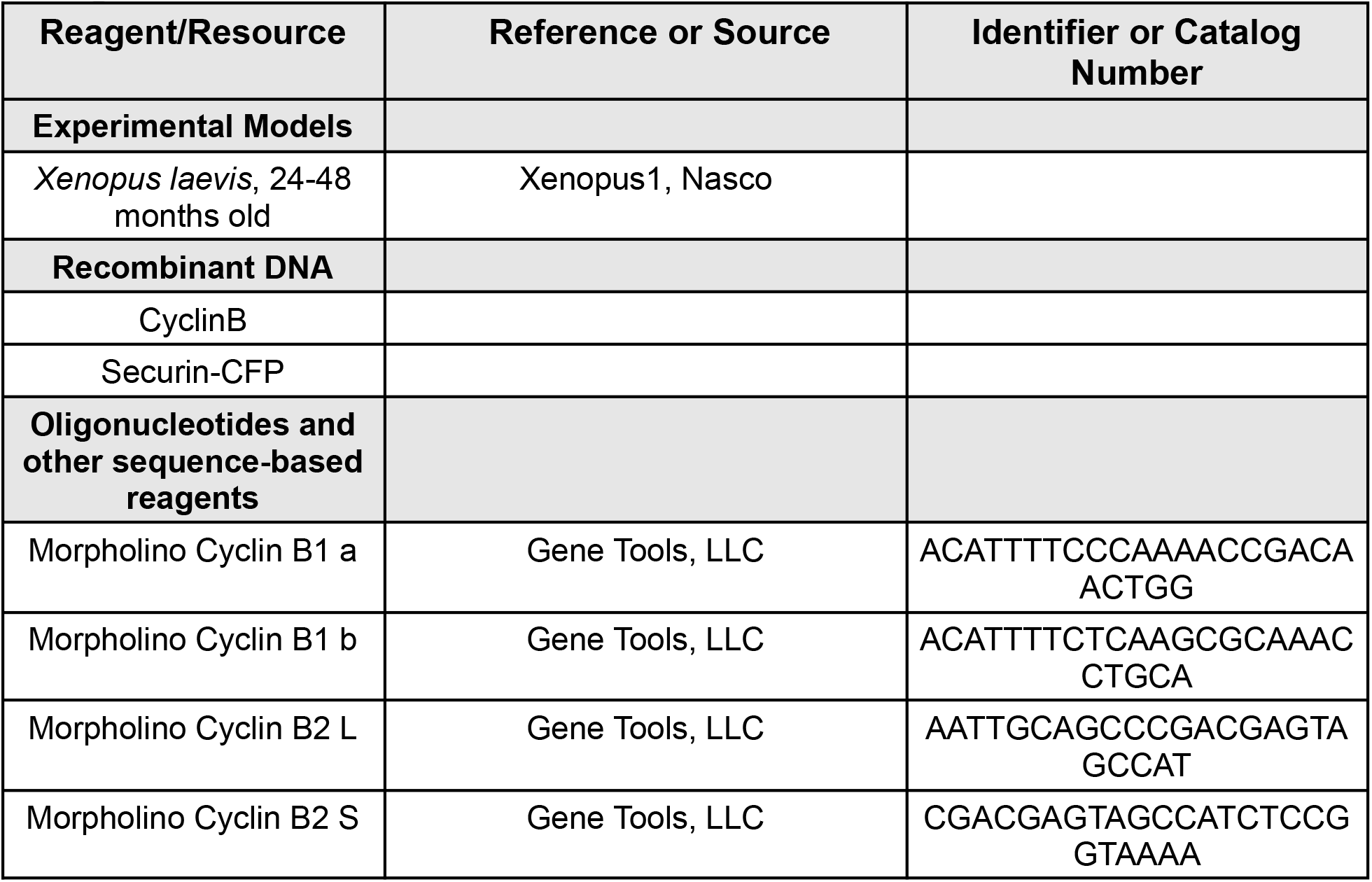
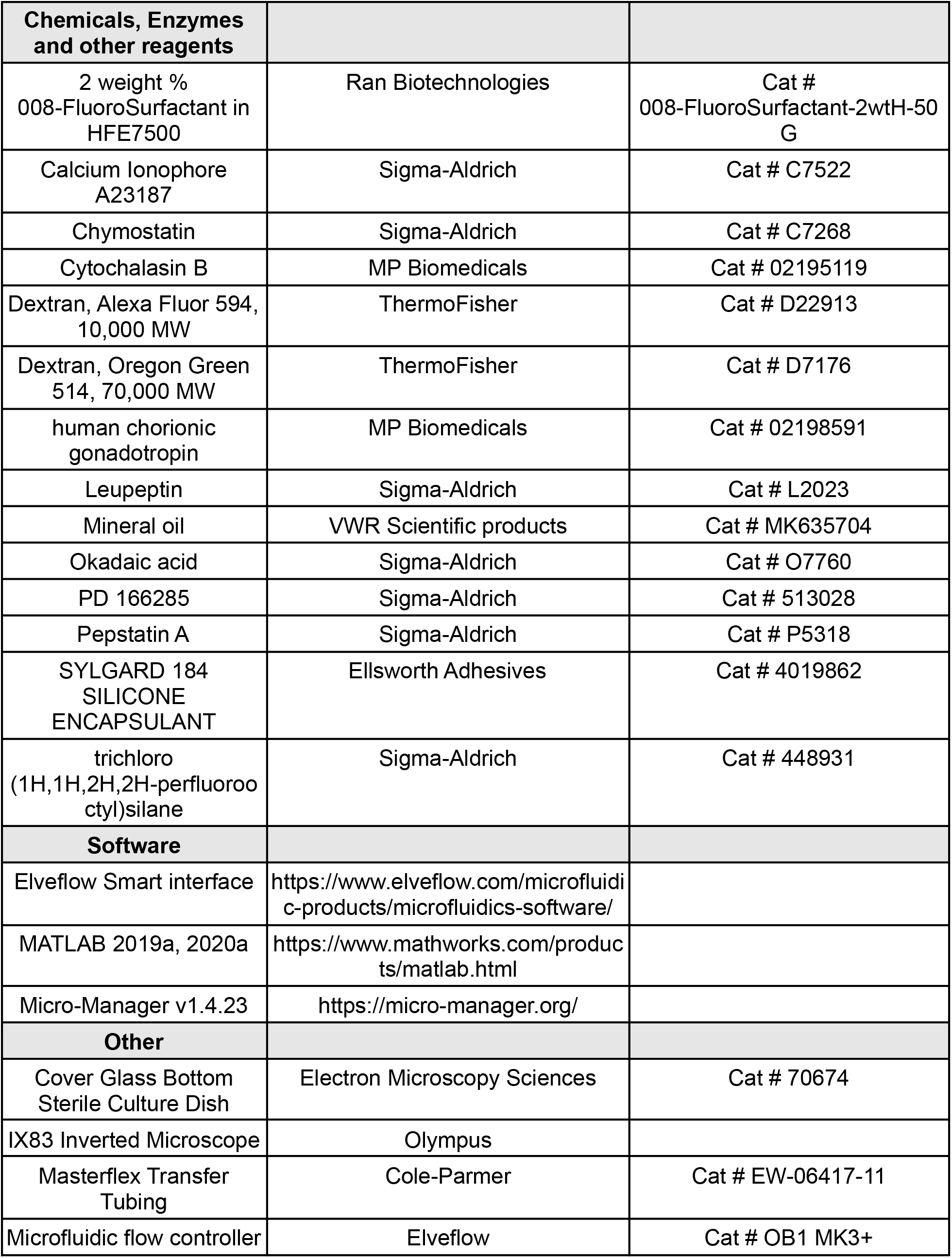
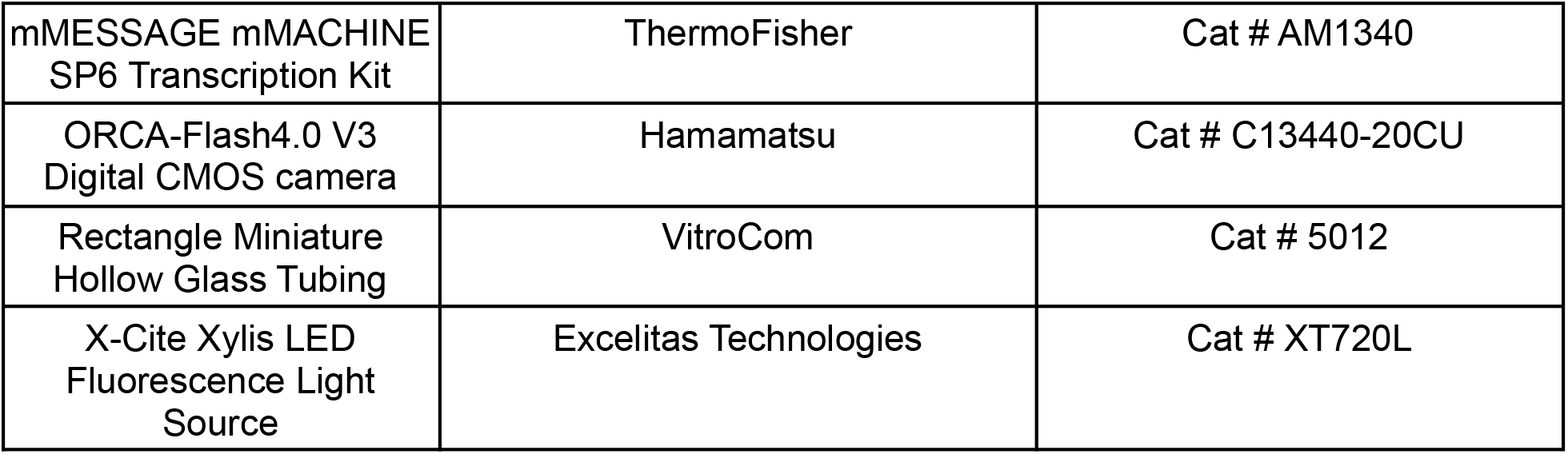

### *Xenopus* cycling extract preparation

The preparation of cycling extract using *Xenopus laevis* eggs has been described in previous publications (Guan et al, 2018a). The frogs are primed with an injection of 100 IU human chorionic gonadotropin (HCG) one week before and again with 600 IU HCG 12-16 hours before harvesting the eggs. The cytosolic materials were extracted via two-step centrifugation at 20,000g. Fluorescent dyes, cell cycle markers, and molecules of interest were then added directly to the extract and gently mixed before droplet generation.

### Droplet generation with microfluidic devices and content tuning

The method for microfluidic device fabrication, droplet generation, and loading were described in our previous report (Sun et al, 2019). Instead of using a syringe pump, we used a multichannel pressure controller (OB1 MK3+, Elveflow, Paris, France) to control the flow with precision. Cycling extracts with different molecules of interest were flown in through different inlets before droplets were generated. The temporal pressure profiles of individual inlets were programmed to change periodically while keeping the combined aqueous pressure constant (Fig S1C, Fig S2F), yielding droplets with different concentrations of these molecules in a wide yet continuous range. The concentrations of the molecules of interest were visualized by co-adding fluorescent dyes of different colors to the extract and creating an index for every droplet, where the fluorescence intensity can be converted to actual concentration when calibrated with reference droplets.

### Time-lapse image acquisition and image analysis

The droplets were loaded into thin glass tubes (inner dimension: 100μm in height, 2mm in width) pre-coated with trichloro(1H,1H,2H,2H-perfluorooctyl)silane to form a 2-D single droplet layer, then immersed in a glass-bottom Petri dish filled with mineral oil to prevent sample evaporation. The dish was then loaded on an inverted fluorescence microscope. Droplets were excited by a LED fluorescence light source (X-Cite Xylis LED Fluorescence Light Source, Excelitas Technologies) and images were then captured by a digital CMOS camera (exposure times: Bright field, 1ms; CFP, 400ms; YFP, 200ms; RFP, 200ms). Automatic stage positioning and imaging were controlled by Micro-Manager, an open-source microscopy software. The images were processed with a custom pipeline of MATLAB scripts. Individual droplets were segmented and tracked using bright-field images. Segmentation was achieved by a watershed algorithm with a seed generated from the Hough circle detection. Tracking was performed by maximizing the segmentation feature correlation between two consecutive time frames. Fluorescent intensity profiles of droplets were then extracted for further analysis.

### Two-ODE model of cell cycle with Cdk1-APC feedback

There are many mathematical models of cell cycle centered on Cdk1-APC feedback with experimentally verified parameters (Novak & Tyson, 1993; Ciliberto et al, 2003; Pomerening et al, 2005; Tsai et al, 2008). Here we use the model presented by Yang and Ferrel (Yang & Ferrell, 2013) for its simplicity. The Cdk1’s two positive feedback with Wee1 and Cdc25 and one negative feedback with APC are incorporated in a two-ODE system. The simplification of feedback could be achieved by assuming fast reaction between a kinase and its substrate. For example, for Cdk1 and Cdc25, we have 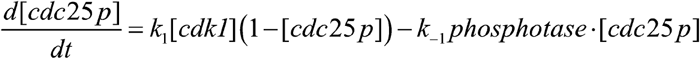, if we assume 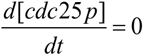 then 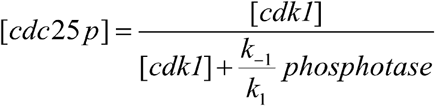. The Hill coefficient is added to the equation to account for experimentally observed hypersensitivity. The final system is shown below

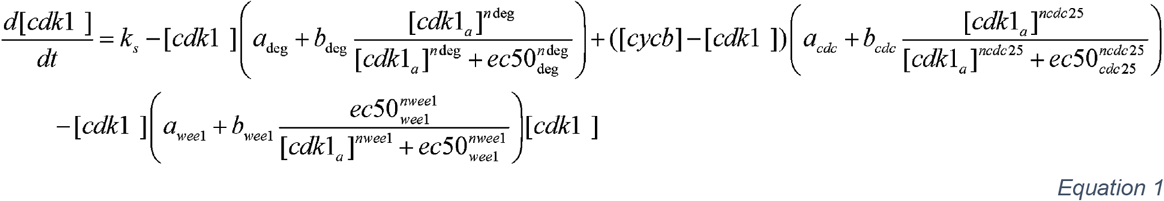

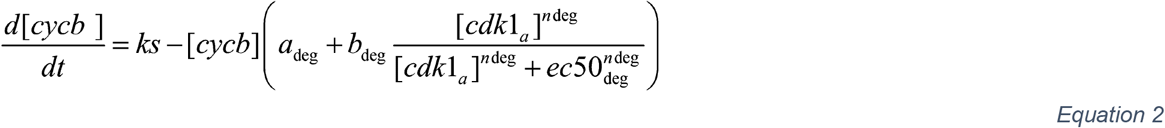

The parameters of this system are shown below. It is slightly changed from the original paper to match experimentally observed periods. The simulation is performed in MatLab 2019b with ode15s method, relative tolerance= 10e-6 and absolute tolerance =10e-8.

### Four-ODE model of cell cycle with Cdk1 and PP2A feedback

The significance of PP2A feedbacks in cell cycle regulation has been under intense investigation, and many models have been proposed to explain the dynamics of Cdk1-PP2A in cell cycle oscillations (Rata et al, 2018; Krasinska et al, 2011; Zhang et al, 2013; Kamenz et al, 2021; Hutter et al, 2017). Here we chose the simplest model that could capture the general dynamics of PP2A feedback with two equations. Combining with the Cdk1-APC system, we get a four-equation system that could explain the general dynamics of the cell cycle. Since here we consider the PP2A to be the phosphatase that counteracts Cdk1, the fast balancing of substrate reaction from the previous model needs to be modified. Take the example of Cdc25, 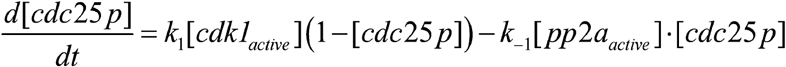. if we assume 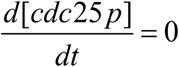, then 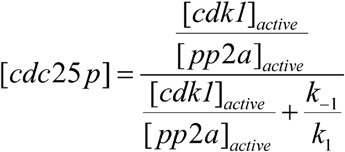. We added Hill coefficient to account for experimentally observed hypersensitivity. Note that the term with Hill coefficient does not represent actual reaction mechanism, and it is just used to describe the experimentally observed responses. After incorporating the static Cdc25, Wee1, and APC activity, we get a four-equation system:

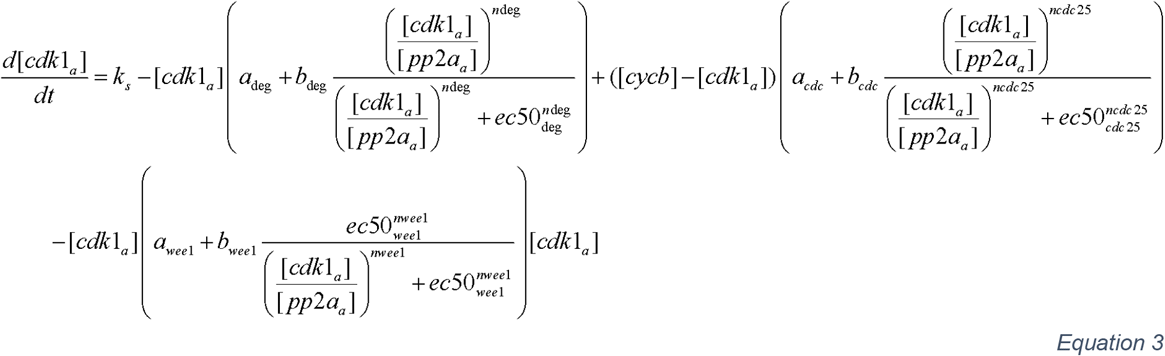

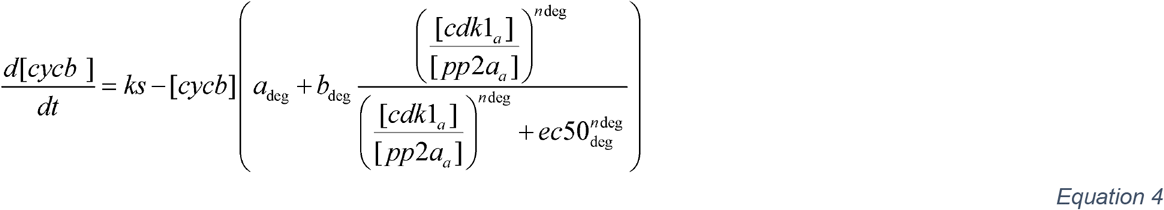

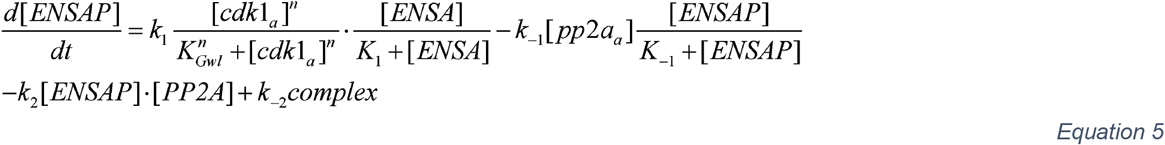

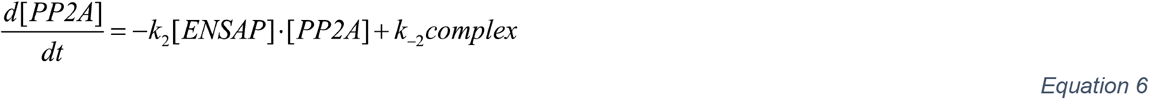

Here the complex represents the PP2A-ENSA complex, note that the PP2A, ENSA, and complex have the following conservations.

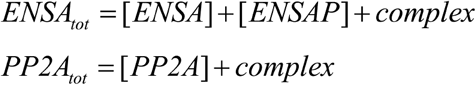

From the observations of previous research, the PP2A may have two different states, and the Cdk1 activity is positively correlated with higher activity PP2A state, so we have active and basal PP2A concentration with the following expression (Kamenz et al, 2021) :

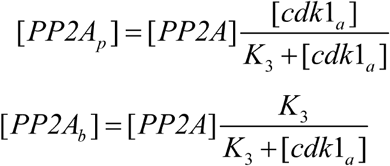

Then we can calculate PP2A activity based on normalized basal and active PP2A concentration

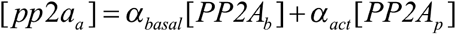

Upon adding OA, the PP2A activity is modified as

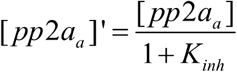

Note that even though Gwl could also be dephosphorylated by PP2A (Williams et al, 2014), previous research has suggested that Gwl is mainly regulated by pp1 (Ma et al, 2016; Rogers et al, 2016; Heim et al, 2015). In addition, hyperphosphorylation is only weakly correlated with Gwl activity (Heim et al, 2015). As a result, the inhibitory effect of PP2A on Gwl is not explicitly indicated in our model.

The simulation is performed in MatLab 2019b with ode15s method, relative tolerance= 10e-6 and absolute tolerance =10e-8. The initial condition is [cdk1_a_]=10, [cycb]=10, [ENSAP]=0.1, [PP2A]=0.1. We use the same parameters as the original model, and the new parameters are chosen to ensure persistent oscillations. The values of the parameters are shown in table 2.

**Table 1.** Parameters for two-ODE systems

| Parameter | Value |
| --- | --- |
| $k_s$ | 1 |
| $a_{deg}$ | 0.012 |
| $b_{deg}$ | 0.02 |
| $a_{cdc}$ | 0.1 |
| $b_{cdc}$ | 0.8 |
| $ec50_{cdc}$ | 35 |
| $n_{cdc}$ | 11 |
| $a_{wee}$ | 0.05 |
| $b_{wee}$ | 0.2 |
| $ec50_{wee}$ | 35 |
| $n_{wee}$ | 3.5 |
| $ec50_{deg}$ | 32 |
| $n_{deg}$ | 17 |

**Table 2.** Parameters for four-ODE systems

| Parameter | Value |
| --- | --- |
| $k_s$ | 1 |
| $a_{deg}$ | 0.01 |
| $b_{deg}$ | 0.04 |
| $a_{cdc}$ | 0.16 |
| $b_{cdc}$ | 0.8 |
| $ec50_{cdc}$ | 35 |
| $n_{cdc}$ | 11 |
| $a_{wee}$ | 0.08 |
| $b_{wee}$ | 0.4 |
| $ec50_{wee}$ | 35 |
| $n_{wee}$ | 3.5 |
| $ec50_{deg}$ | 32 |
| $n_{deg}$ | 17 |
| $k_1$ | 15 |
| $K_1$ | 0.1 |
| $k_{-1}$ | 1.5 |
| $K_{-1}$ | 0.01 |
| $k_2$ | 600 |
| $k_{-2}$ | 15 |
| $n$ | 5 |
| $K_3$ | 10 |
| $a_{basal}$ | 0.024 |
| $K_{Gwl}$ | 100 |
| $\alpha_{act}$ | 1.2 |
| $PP2A_{tot}$ | 1 |
| $ENSA_{tot}$ | 2 |

## Acknowledgements

This work was supported by NSF (MCB #1817909; Early Career #1553031), NIH (NIGMS #R35GM119688), and Alfred P. Sloan Foundation.

## Conflict of interest

The authors declare no conflict of interest.

## Appendix

**Appendix Figure S1.**
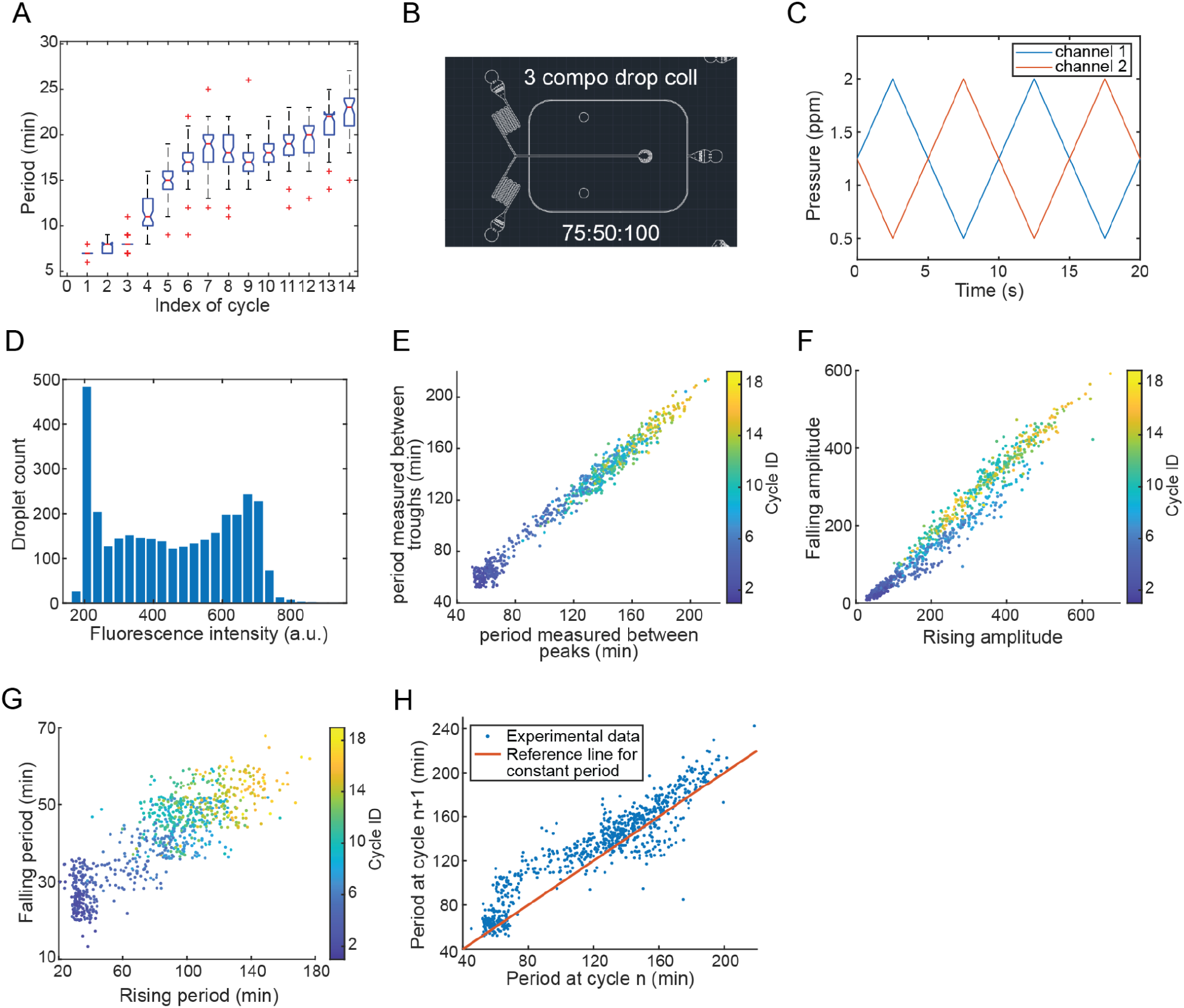
**A** Oscillation period of each mitotic cycle in droplets. The extracts used were not manipulated except the addition of securin-CFP and energy mix. **B** Design of 2-channel microfluidic device for droplet generation. Extracts were flown in from the 2 inlets on the left, surfactant oil was flown in from the right, droplets were collected in a well in the middle punched in after fabrication. **C** Temporal pressure profile of the 2 extract channels for continuous content tuning. The sum of total pressure is kept constant. **D** Histogram of fluorescent dye intensity in droplets generated by 2-channel Cyclin B mRNA tuning. The peaks at both ends were caused by intentionally generating more droplets at extreme conditions to ensure the tuning covered the whole desired range of concentration. **E** Comparison between oscillation period calculated between peaks and troughs. Color bar indicates cycle ID. **F** Comparison between rising amplitude and falling amplitude of the securin-CFP signal of each cycle. Color bar indicates cycle ID. **G** Comparison between periods of rising phase and falling phase of each cycle. Color bar indicates cycle ID. **H** Comparison between 2 consecutive cycles.

**Appendix Figure S2.**
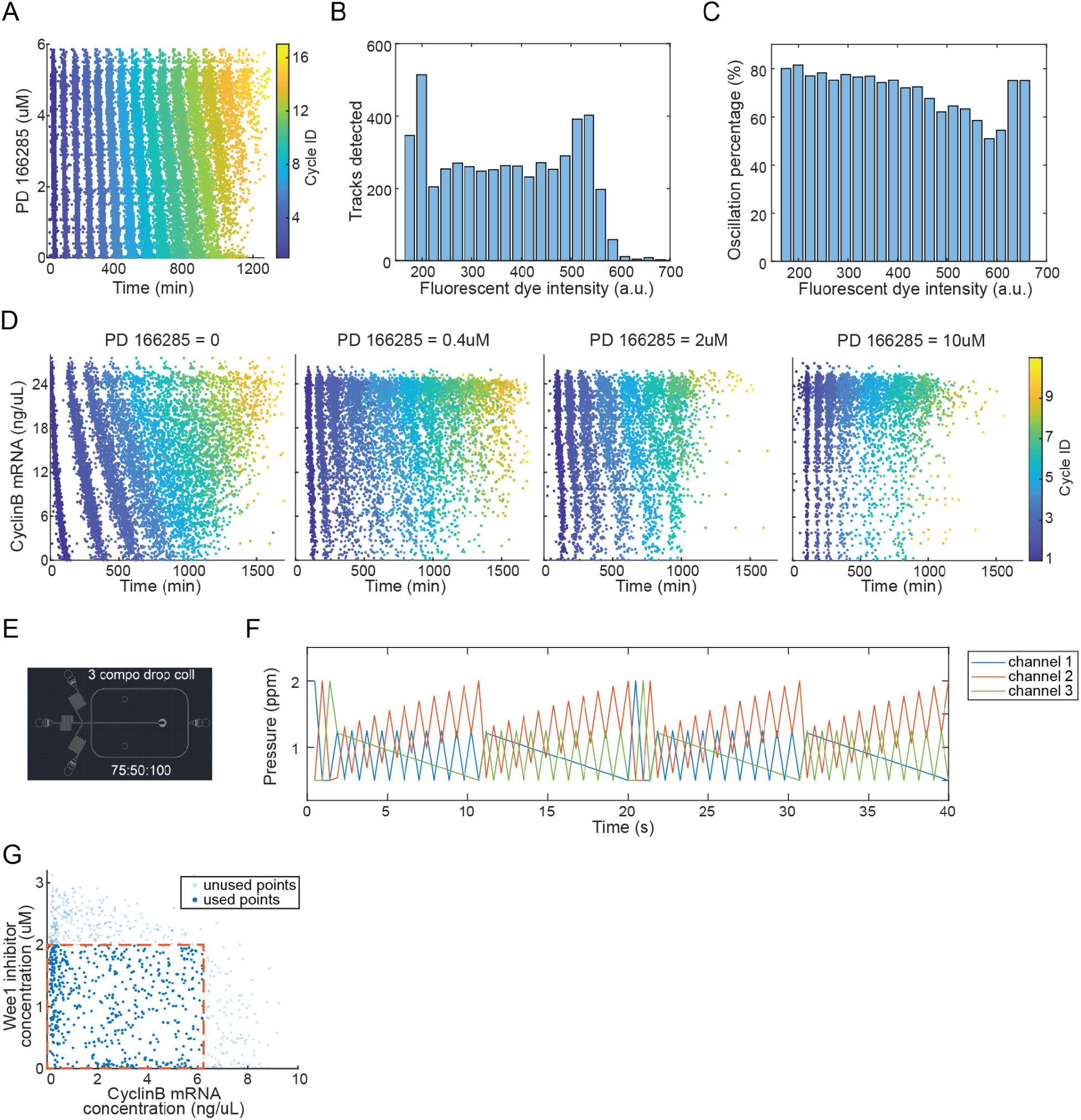
**A** Raster plot of mitotic oscillations of droplets with different PD 166285 concentrations. Each point represents one peak in the securin-CFP signal of one droplet. The droplets were generated by 2-channel tuning to change PD 166285 concentration. Color bar indicates cycle ID. **B** Histogram of fluorescent dye intensity in droplets generated by 2-channel PD166285 tuning. **C** Percentage of oscillation droplets at different PD 166285 concentrations, indicated by fluorescent dye intensities. **D** Raster plot of mitotic oscillations of droplets with different cyclin B mRNA concentrations at 4 separate PD 166285 concentrations (0, 0.4, 2, 10µM). Color bar indicates cycle ID. **E** Design of 3-channel microfluidic device for droplet generation. Extracts were flown in from the 3 inlets on the left, surfactant oil was flown in from the right, droplets were collected in a well in the middle punched in after fabrication. **F** Temporal pressure profile of the 3 extract channels for continuous content tuning. The sum of total pressure is kept constant. The cycles are designed so the parameter space is evenly scanned. **G** The resulting concentration of Cyclin B mRNA and PD 166285 in droplets generated by 3-channel tuning. Only data points in the selected area were used for further analyses.

**Appendix Figure S3.**
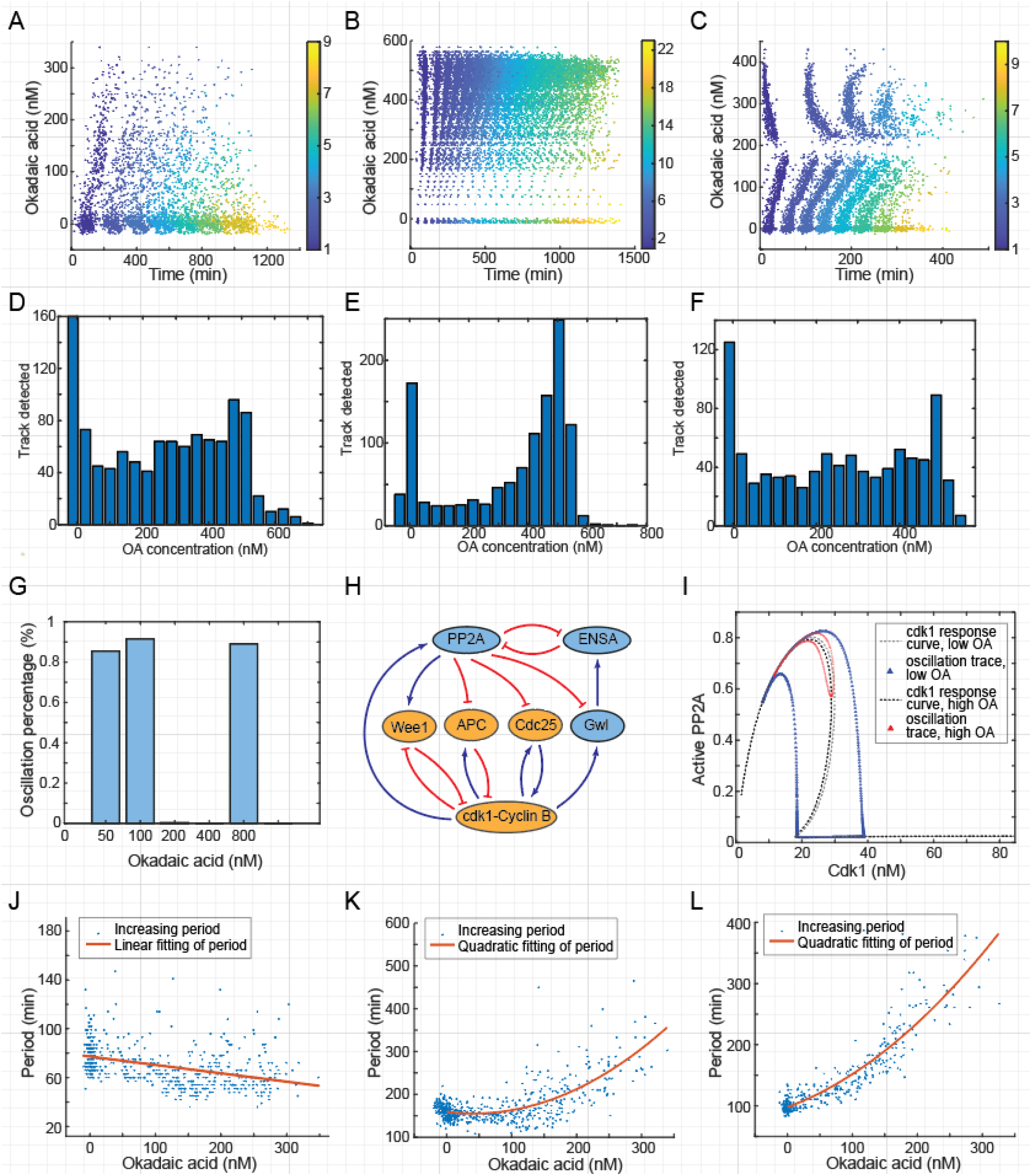
**A, B, C** Representative raster plot of type 1 (A) 2 (B) 3 (C) OA response. Droplets are sorted by OA concentration, and the color codes the peak index. **D, E, F** Histogram of tracked droplets corresponding to type 1 (D) 2 (E) 3 (F) OA response. **G** Percentage of droplets that show oscillatory behavior when adding different concentrations of OA. Note that no oscillator is detected in 200 nM or 800 nM conditions. **H** Interaction diagram of PP2A model. Red circuits indicate the mitotic entry positive feedback, and the blue shows mitotic exit positive feedback. **I** Simulated limit cycle on the Cdk1-PP2A phase plane. The dotted line shows active PP2A response curve to Cdk1 activity when type 3 oscillation presents. Blue and red curves show low OA and medium OA conditions respectively. **J, K, L** Oscillation period changes when tuning OA. Showing three different responses. monotonic decreasing (J), decreasing and then increasing (K), and monotonic increasing(L). Red lines are fitted curves with linear (J) or quadratic models (K, L).

## Notes

### Competing Interest Statement

The authors have declared no competing interest.

